# Predicting drowning from sea and weather forecasts: development and validation of a model on surf beaches of southwestern France

**DOI:** 10.1101/721142

**Authors:** Éric Tellier, Bruno Simonnet, Cédric Gil-Jardiné, Marion Bailhache, Bruno Castelle, L. Rachid Salmi

**Author notes:** **Corresponding author:** Éric Tellier.

## Abstract

**Objective:** To predict the risk of drowning along the surf beaches of Gironde, southwestern France.

**Methods:** Data on rescues and drownings were collected from the Medical Emergency Center of Gironde (SAMU 33). Seasonality, holidays, weekends, weather, and sea conditions were considered potentially predictive. Logistic regression models were fitted with data from 2011–2013 and used to predict 2015–2017 events employing weather and ocean forecasts.

**Results:** Air temperature, wave parameters, seasonality, and holidays were associated with drownings. Prospective validation was performed on 617 days, covering 232 events (rescues and drownings) reported on 104 different days. The area under the curve (AUC) of the daily risk prediction model (combined with 3-day forecasts) was 0.82 [95% confidence interval (95% CI) 0.79−0.86]. The AUC of the 3-hour step model was 0.85 (95% CI 0.81−0.88).

**Conclusions:** Drowning events along the Gironde surf coast can be anticipated up to 3 days in advance.Preventative messages and rescue preparations could be increased as the forecast risk increased, especially during the off-peak season, when the number of available rescuers is low.

## INTRODUCTION

According to the 2017 Global Burden of Disease study, drowning (294,000 fatalities yearly) is a major cause of non-intentional deaths from injury worldwide^1^. In France, the national public health agency (Santé Publique France) performs a national study every 3 years, registering all cases of drowning leading to hospitalization or death between June 1 and September 15. In 2015, there were 1,266 drownings in France, with 637 (50.3%) occurring along the seashore^2^. In a previous study on the surf beaches of Gironde, southwestern France, 576 people required rescue over 6 years; there were 24 fatalities^3^. In terms of the length of the coastline, the annual mean was 3.3 deaths/100 km, a rate comparable to the highest recorded along the US coastline^4^.

The Gironde coast is a 126-km-long stretch of sandy beaches exposed to high-energy waves that drive intense, narrow seaward-flowing jets of water termed “rip currents”; these flow through deep channels in sandbars that parallel the shore. In southwest France, rip-current activity normal to the shore increases with increasing wave height and period, lower tide height, and variability in beach morphology^5^. A previous study showed^6^ that these currents cause 79% of drownings. Rip currents are the leading causes of rescues and drownings off many surf coasts worldwide^4,7–10^.

Drowning is sudden; prevention is key when the aim is to reduce the incidence of drowning^11–13^. Primary prevention may modify beachgoer behavior^14^. If a drowning appears imminent, a fast response by paramedics (and a medical team if necessary) is essential^15^. Lifeguards can impart preventative messages^16^, reducing the need for medical attention and cardiopulmonary resuscitation of drowning victims^17,18^.

Drowning prevention on the Gironde beaches features patrolled areas, signs at most beach entrances, and leaflets describing the rip current and shore-break hazards. However, the beaches are not patrolled during the entire bathing season, which extends from April to October. Most lifeguard stations are open only in July and August; the locations most frequented by tourists are patrolled from mid-June to mid-September. On weekends in May and June, some areas are watched, depending on local authorities. The mayor is responsible for beach supervision, which is regionally coordinated by the departmental prefect in collaboration with the pre-hospital care department of Bordeaux University Hospital. During high season, rescue helicopters are on standby. On low-season weekends, one helicopter may be on duty, depending on the regional authority.

Models predicting the risk of drowning would be useful if they enhanced the preventative measures taken to reduce risk. Predictive models of rip currents have been implemented in Florida, Puerto Rico^19^, Mexico^20^, India^9^, and Great Britain^21^. The models are based on physical hazards, and have been evaluated both retrospectively and in the field. To the best of our knowledge, they have not been prospectively evaluated; forecasts have not been compared to actual drownings.

Exposure to a rip hazard increases as the number of swimmers rises. Attendance rises on holidays, weekends, and with increased air temperature and less cloud cover (nebulosity); the number of bathers reflects air and water temperatures and (possibly) wind speed. As the risk of drowning is a combination of the hazard per se and exposure to it, and as the latter is poorly quantified, we created a model including parameters reflecting exposure to rip currents. We assessed whether drownings off Gironde beaches could be anticipated using a risk prediction model based on forecast weather and ocean conditions.

## MATERIALS AND METHODS

### Study setting

We performed an observational study along the French Atlantic coastline of Gironde. We first developed a model based on medical emergency calls from beaches, along with weather and sea conditions, in 2011, 2012, and 2013. We evaluated only the bathing season (April to October). We used the model to assess whether weather forecasts accurately predicted events that occurred from April to October in 2015, 2016, and 2017. We used the RiGoR guidelines to address common sources of bias in risk-prediction models,^22^ and we adhered to the STROBE statement for observational studies^23^.

### Data sources

#### Medical emergency calls

In Gironde, every medical emergency call from a beachgoer or lifeguard is received by a single medical emergency call center (SAMU, Service d’Aide Médicale d’Urgence). During each call, a physician records all information given by the caller, paramedics, and (when applicable) pre-hospital care teams. All calls dealing with rescue from water or drowning were included in the data for this study; these were the events of interest. “Rescue” refers to a need for evacuation from the water, and “drowning” refers to respiratory impairment caused by submersion or immersion, as defined by the World Health Organization^24^. We excluded calls lacking victims, training calls, and duplicates. As every instance of a need for medical advice or a pre-hospital care team triggered a call, we considered that all events of importance would be identified. Information on every call was carefully read to avoid errors. Intentional drownings and drownings associated with known diseases were excluded.

#### Environmental conditions

Hourly tidal data were modeled by the “Service Hydrographique et Océanographique de la Marine” (SHOM, authorization no. 296/2014) using the Lacanau shore as the reference. Lacanau is located in the approximate center of the study area; according to the SHOM, the maximum tide phase lag over the entire study area is approximately 15 min. Wave conditions were measured every 30 min by the CANDHIS buoy^25^ located at 044°39.150’N and 001°26.800’W. The wave propagation time from the buoy to the coast is about 1 h. Observed and forecast meteorological and wave conditions were provided by Météo-France, the French national meteorological service. We used data from Cap Ferret; Météo-France claims that these well-represent the weather along the entire Gironde coast. Forecast data were available for up to 3 days and at 3-h steps (7:00 am UTC, 10:00 am, etc.). We recorded sea height, the wave factor (the product of wave height and period), and the wave incidence factor (as defined in Appendix 1). We also recorded wind speed and direction, air and water temperatures, and nebulosity. Other factors influencing beach attendance were the season and type of day. High season was defined as the period from June 15 to September 15, when most lifeguard stations are open. We distinguished between weekdays, weekends, and holidays.

#### Statistical methods

We fitted two logistic regression models: a “daily model” predicting the overall risk of at least one drowning on a given day, and a “3-h-step model” predicting the risks at different times of the day (9:30 am−12:29 pm, 12:30−3:29 pm, 3:30−6:29 pm, and 6:30-9:30 pm; all local times). Given differences in the data collection modes between the training and validation periods, we checked data coherence both visually and using the Wilcoxon–Mann–Whitney and Student’s t- tests.

Days for which data were lacking were removed from the analysis. Prospective cohort data (including variable selection) were not used during model development. We transformed the wave parameters (Appendix 1). We categorized non-log-linear quantitative variables; these were first divided into quantiles and then reduced using the Akaike Information Criterion (AIC) in a multivariate context^26^. Model selection used the AIC to perform interaction checks; we tested all possible models^27^. Odds ratios (ORs) with 95% confidence intervals (CIs) were computed as bootstrap estimates. We checked that residual autocorrelation was absent. Goodness of fit was assessed using the Le Cessie-Van Houwelingen test^28^. Calibration was assessed graphically employing a locally weighted, least-square regression smoother^29^ and the Spiegelhalter Z-test. Discriminatory power was assessed using receiver operator characteristic (ROC) curves based in data from each cohort. Fit and validation accuracies were assessed via Brier scoring. The relative importance of the selected predictors was assessed by deriving the associated chi-squared proportions. The outcomes derived using 1-, 2-, and 3-day forecasts were compared by drawing ROC curves using the Delong and Venkatraman method for paired data;^30–32^ we applied Holm– Bonferroni corrections. We created a five-level risk scale using the quintiles of the fitted probabilities. All analyses employed R software^33^ running the RMS^29^ and pROC packages^32^.

#### Ethics

Data collection was approved by the French national committee protecting data privacy (Commission Nationale de l’Informatique et des Libertés, CNIL), provided that only compiled (anonymized) data would be published. French law states that a retrospective observational study does not require ethics committee authorization.

## RESULTS

Retrospective data were lacking for 77 days because of a buoy failure, and for 26 prospective days (21 because of data-link loss and 5 because of server unavailability). We analyzed 563 days during 2011–2013; 242 rescues and drownings were reported on 108 different days. In 2015–2017, data were available for 612 days; there were 232 events on 104 different days (Table 1). All retrospective and prospective cohort data were consistent, except for wind speed, which differed significantly between prospective and retrospective data, and nebulosity, which was measured by different means over the retrospective and prospective periods. Both were excluded from prospective analyses.

**Table 1.** Description of days without and with rescues and/or drownings. Meteorological and wave conditions (medians and quartiles) and the characteristics of days on which rescues and/or drownings occurred along the Gironde coast of southwestern France.

|  | 2011–2013 |  |  |  | 2015–2017 |  |  |  |
| --- | --- | --- | --- | --- | --- | --- | --- | --- |
|  | Days without events<sup>a</sup> (n = 455) |  | Days with events (n = 108) |  | Days without events (n = 513) |  | Days with events (n = 104) |  |
| Wave factor<sup>b</sup>, m × s | 11.3 | 5.6–14.6 | 10.3 | 6.5–12.9 | 11.3 | 5.4–14.4 | 14.5 | 8.0–18.0 |
| Wave angle factor<sup>c</sup> | 0.80 | 0.69–0.99 | 0.89 | 0.85–0.99 | 0.80 | 0.74–0.97 | 0.88 | 0.83–0.97 |
| Nebulosity (0–4)<sup>d</sup> | 2.8 | 2.0–3.7 | 2.4 | 1.5–3.0 |  | – |  | – |
| Air temperature, °C | 21.6 | 19.2–23.9 | 25.2 | 23.3–26.6 | 21.5 | 19.0–24.0 | 25.5 | 23.0–27.0 |
| Water temperature, °C | 19.0 | 17.6–20.9 | 21.3 | 20.3–22.5 | 18.0 | 16.0–20.0 | 20.3 | 20.0–21.0 |
| Wind speed<sup>e</sup>, m/s | 7.0 | 5.3–8.2 | 6.6 | 5.3–7.3 | 4.9 | 2.7–5.5 | 4.2 | 2.7–5.5 |
| Season<sup>f</sup>, n (%) |  |  |  |  |  |  |  |  |
| High | 187 | (41.1) | 90 | (83.3) | 199 | (38.8) | 79 | (76.0) |
| Low | 268 | (58.9) | 18 | (16.7) | 314 | (61.2) | 25 | (24.0) |
| Type of day, n (%) |  |  |  |  |  |  |  |  |
| Weekday | 213 | (46.8) | 18 | (16.7) | 233 | (45.4) | 24 | (23.1) |
| Weekend | 82 | (18.0) | 12 | (11.1) | 88 | (17.2) | 17 | (16.3) |
| Vacation | 160 | (35.1) | 78 | (72.2) | 192 | (37.4) | 63 | (60.6) |

<sup>a</sup> Events include rescues and drownings.
<sup>b</sup> Wave factor: wave height (m) times wave period (s).
<sup>c</sup> The wave angle factor ranges from 0 to 1; see Appendix 1 for details.
<sup>d</sup> Forecast values not shown because of differences in the modes of data measurement.
<sup>e</sup> Significant differences between observed and forecast data.
<sup>f</sup> High season: June 15 to September 15.

The final, predictive, daily risk model included wave and wave angle factors, air temperature, type of day, and season (Table 2). The model predicting risk at 3-h steps featured sea height, wave parameters, air temperature, time of day, type of day, and season (Table 3). Variation in the daily model was attributable principally to air temperature (proportion of the overall chi-squared value, 40.9%), wave factors (21.7%), and time of day (16.2%). The principal 3-h-step model predictors were air temperature (28.9%), the time of day (17.8%), and wave factors (12.6%).

**Table 2:** Factors associated with daily rescues and drownings along the Gironde coast. Univariate and multivariate analyses performed with the aid of logistic regression models using retrospective data from 2011–2013.

| | Crude OR | (95% CI) | Adj. OR | (95% CI) | $\chi^2$ |
| --- | --- | --- | --- | --- | --- |
| Wave factor<sup>a</sup>, m × s |  |  |  |  | 22.8 |
| <5.2 |  | Ref. |  | Ref. |  |
| 5.2–9.2 | 3.89 | (2.03, 9.81) | 6.41 | (2.93, 18.3) |  |
| >9.2 | 1.96 | (1.01, 4.50) | 7.10 | (3.09, 22.7) |  |
| Wave angle factor<sup>a</sup> | 1.83 | (1.40, 2.62) | 2.27 | (1.58, 3.70) | 13.2 |
| Nebulosity<sup>b,c</sup> | 0.52 | (0.37, 0.74) | — |  |  |
| Air temperature<sup>a</sup>, °C |  |  |  |  | 43.0 |
| ≤21 |  | Ref. |  | Ref. |  |
| 21–23.5 | 6.58 | (3.03, 19.2) | 4.79 | (1.93, 16.6) |  |
| >23.5 | 19.10 | (9.62, 61.8) | 12.20 | (4.69, 52.3) |  |
| Water temperature<sup>a</sup>, °C |  |  |  |  |  |
| ≤19.5 |  | Ref. |  | — |  |
| 19.5–21.3 | 3.95 | (2.13, 9.14) |  |  |  |
| >21.3 | 13.18 | (7.60, 27.0) |  |  |  |
| Wind speed<sup>b</sup>, m/s |  |  |  |  |  |
| ≤4.3 |  | Ref. |  | — |  |
| 4.3–6.3 | 1.71 | (1.00, 3.14) |  |  |  |
| >6.3 | 0.84 | (0.45, 1.66) |  |  |  |
| Season |  |  |  |  | 10.4 |
| Low |  | Ref. |  | Ref. |  |
| High | 7.17 | (4.38, 13.5) | 3.98 | (1.44, 6.11) |  |
| Type of day |  |  |  |  | 17.1 |
| Weekday |  | Ref. |  | Ref. |  |
| Weekend | 1.73 | (0.71, 3.62) | 2.96 | (1.06, 7.98) |  |
| Holidays | 5.77 | (3.49, 10.8) | 4.25 | (2.19, 9.75) |  |

*Note.* OR = odds ratio, CI = confidence interval
<sup>a</sup> Daily maximal value.
<sup>b</sup> Daily mean value.
<sup>c</sup> Not incorporated into multivariate analyses because of differences in measurement modes.

**Table 3:**
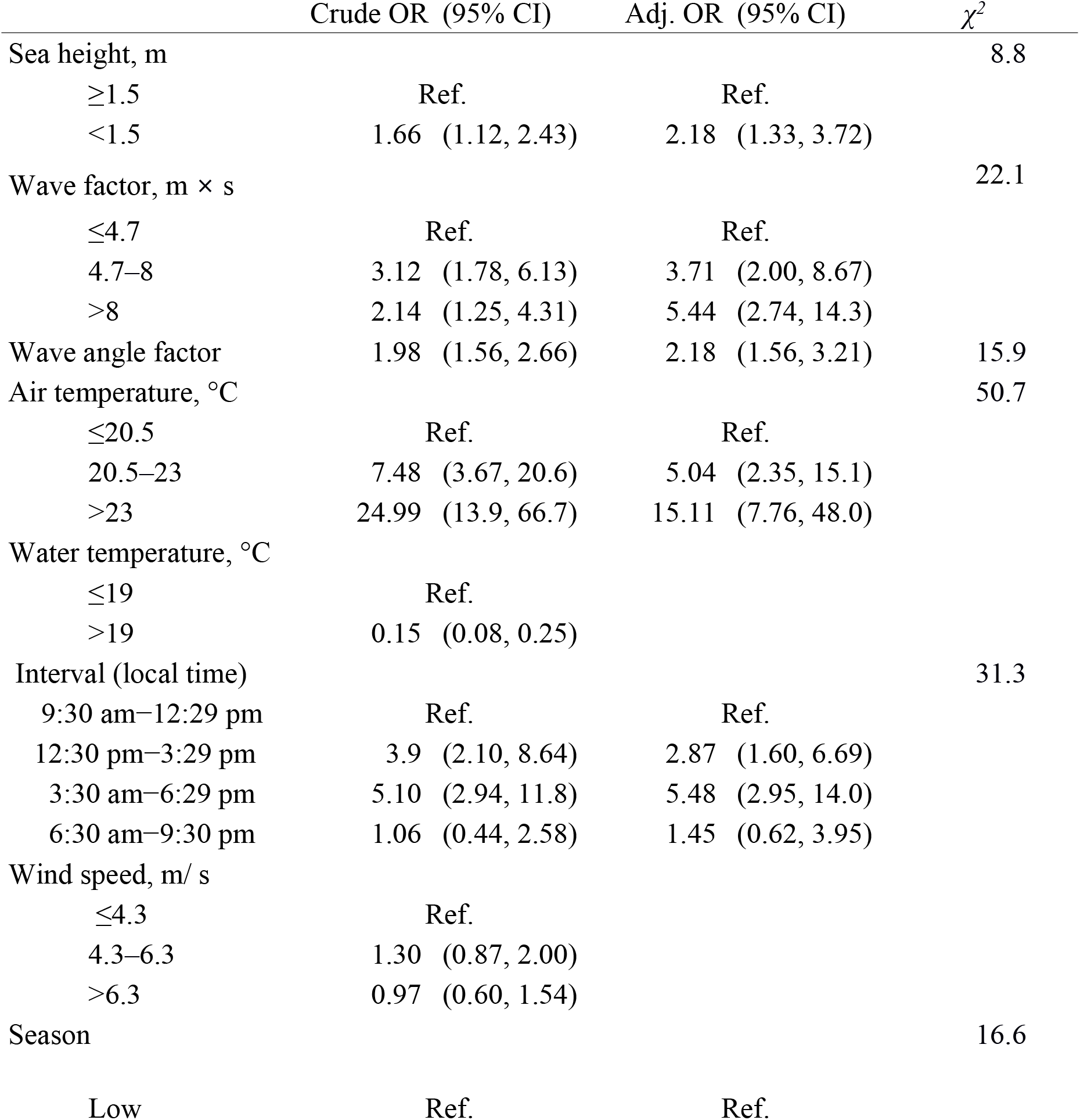
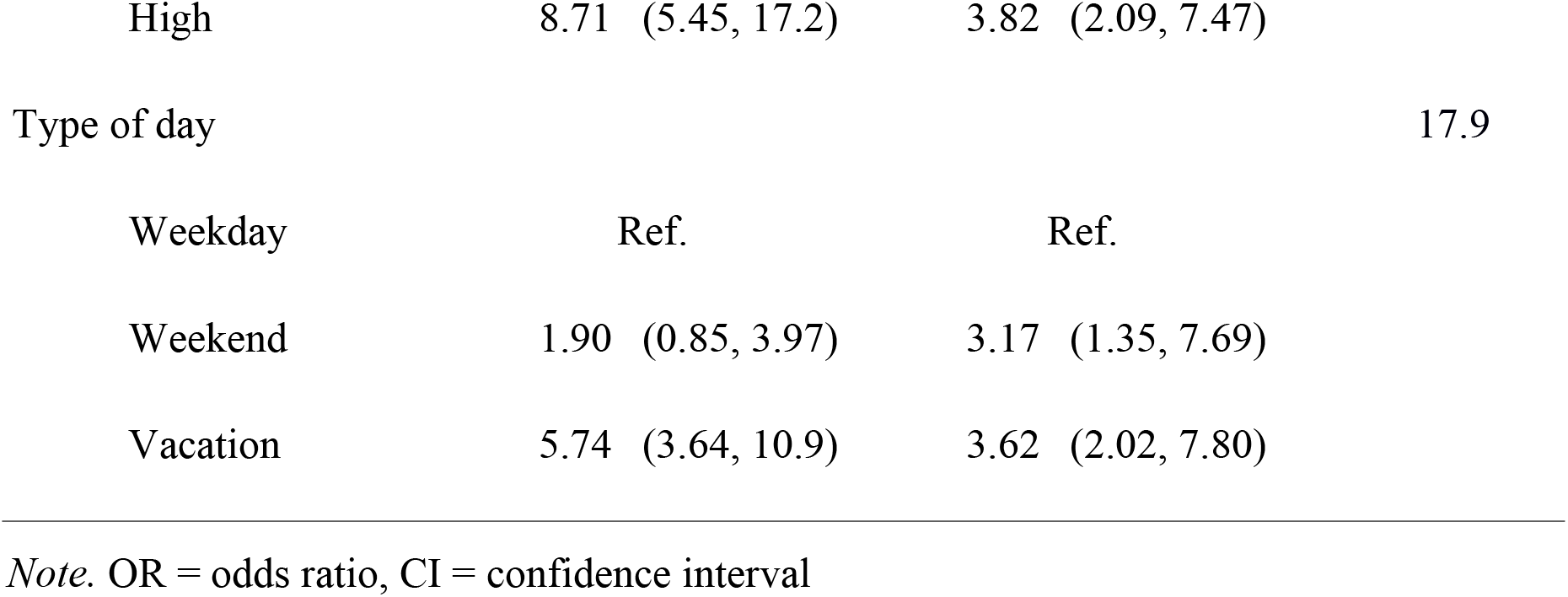
Factors associated with rescues and drownings during 3-h periods. Univariate and multivariate analyses performed with the aid of logistic regression models for risk of drowning along the Gironde coast during 3-h periods using retrospective data from 2011–2013.

The daily model had areas under the curves (AUCs) of 0.88 (95% CI 0.84–0.91) for 2011–2013 and 0.82 (95% CI 0.78–0.86) for 2015–2017. The 3-h risk model had AUCs of 0.89 (95% CI 0.87–0.92) for 2011–2013 and 0.85 (95% CI 0.81–0.88) for 2015–2017. Model outcomes did not differ when forecasts for 1, 2, and 3 days were used (*p* > 0.05). Both models were well calibrated in terms of retrospective data (goodness-of-fit test *p* = 0.20 for the daily model, *p* = 0.53 for the 3-h-step model). Both models exhibited significant *p*-values on Spiegelhalter Z-testing of prospective data, evidencing a lack of calibration: the daily model tended to over-predict days with risks of drowning >0.5; the 3-h step model over-predicted risks as low as 0.1.

Using prospective data, we found that assessment of the drowning risk using the five-level scale missed 1 of 158 days featuring a rescue at the lowest risk level (0.6%). The missed case was a rescued male who presented without a cough and was discharged on site. The prospective data predicted 45.8% of days with rescue events at the highest risk level (Table 4). The 3-h step model missed 2 of 481 rescues, one at the lowest level (0.4%) and one at the highest (15.7%).

**Table 4.** Observed rescues vs. predicted drowning risk. Observed rescues and drownings by predicted risk level derived using regression models exploiting 3-day forecasts; Gironde, southwestern France.

| Risk Level | 2011–2013 |  |  | 2015–2017 |  |  |
| --- | --- | --- | --- | --- | --- | --- |
|  | Events<sup>a</sup> | Total | % | Events | Total | % |
| Daily model<sup>b</sup> |  |  |  |  |  |  |
| 1 | 3 | 113 | 2.7 | 1 | 172 | 0.6 |
| 2 | 1 | 112 | 0.9 | 8 | 105 | 7.1 |
| 3 | 11 | 113 | 9.7 | 15 | 95 | 15.8 |
| 4 | 23 | 112 | 20.5 | 23 | 88 | 26.1 |
| 5 | 70 | 113 | 61.9 | 66 | 144 | 45.8 |
| 3-h-step model<sup>c</sup> |  |  |  |  |  |  |
| 1 | 0 | 395 | 0.0 | 2 | 481 | 0.4 |
| 2 | 3 | 394 | 0.8 | 3 | 381 | 0.8 |
| 3 | 6 | 394 | 1.5 | 10 | 309 | 3.2 |
| 4 | 19 | 394 | 4.8 | 22 | 330 | 6.7 |
| 5 | 103 | 394 | 26.1 | 58 | 369 | 15.7 |

<sup>a</sup> Events: Rescues and drownings.
<sup>b</sup> Goodness of fit: $p = 0.20$ , calibration test $p < 0.001$ .
Brier scores: 0.10 for 2011–2013, 0.12 for 2015–2017.
<sup>c</sup> Goodness of fit: $p = 0.53$ , calibration test $p = 0.10$ .
Brier scores: 0.05 for 2011–2013, 0.05 for 2015–2017.

The missed case at the lowest risk level in the retrospective cohort occurred during moderate wave conditions (wave factor ~8 m × s) and at low wave incidence (0.47), but the victim required only rescue, was asymptomatic upon rescue, and was not evacuated. The second missed event occurred at the tip of the Cap Ferret sandspit, which lacks wave-driven rip currents. The last missed event occurred at La Salie Nord, adjacent to (south of) the Arcachon inlet, under moderate wave conditions.

The 3-h step model missed two events in the prospective cohort, both in September 2017, and both occurring under strong wave conditions (wave factor >15 m × s). One was a surfer; the activity pursued by the other victim was not recorded. Both cases were minor and were treated in the local hospital.

## DISCUSSION

This study is the first to analyze the association of forecast sea and weather conditions with drowning risk. Air temperature, wave angle, and the wave factor were the primary environmental predictors; the type of day and the season were also significant, but less important, predictors. Given the availability of extensive physical data on rip currents, we were able to build a tool that accurately predicted drowning risk.

Warm weather increases sea exposure and therefore the risk of drowning, consistent with other studies^34,35^. Wave parameters influencing rip current flow velocity were significant predictors of drowning, consistent with the results of physical models^21^. As the wind parameters differed between the retrospective and prospective periods, we could not use these parameters, although they might have further improved the models. Nebulosity was measured differently during the two periods and thus could not be incorporated into the models. Although univariate analysis showed that nebulosity was a significant predictor of drowning, it is strongly correlated with air temperature. Future models should integrate predicted rather than observed measures.

Our models tend to overestimate the risk on days associated with moderate to high risks; some variables may thus be unknown. First, the summer beach morphology along the Gironde coast is very variable; beach slopes and channel depths differ markedly, and the rip current hazard changes from one summer to the next. Moreover, drownings are certainly under-reported to the Emergency Call Center; reporting rates may vary over time. We could not directly estimate exposure, as beach attendance is not measured in Gironde.

Turning to the missed events, two occurred in sectors adjacent to the Arcachon lagoon inlet, where local, strong tide-driven currents develop at low tide, constituting a major hazard. We hypothesize that the missed events were attributable to these currents. This highlights the need to carefully target preventative messages; the primary hazards vary locally.

Use of the 1-, 2-, and 3-day forecasts yielded similar results; this will aid in the efficient deployment of lifeguards and rescue equipment. Accurate local forecasts more than 3 days ahead are not available.

How may our findings save lives? The use of a binary scale would trigger many false alarms; thus, we considered that a five-level scale was more appropriate, as such scales are used to predict other risks posed by natural hazards (such as snow avalanches). Our scale should be improved using a risk utility function, which remains to be specifically determined. We concede that our present levels are arbitrary; we must still explore what beachgoers and decision-makers consider to be “low,” “moderate,” or “acceptable” risks.

Any message suggested by our models must be consistent with “messages” imparted by beach flags. These flags can be “green” (no or minor hazard, bathing supervised), “yellow” (hazard, bathing supervised) or “red” (major hazard, no bathing allowed). They are determined by lifeguards based on wave conditions, water temperature, and beach attendance, and may vary depending (for example) on lifeguard experience. Unpublished local reports indicate that green and red flags are rarely raised during the high season on the Gironde coast.

We are confident that our model can be adapted to similar beaches with rip currents, but complete generalization of our findings is inappropriate in the absence of more data. Lifeguard knowledge and the physical parameters of natural hazards require attention. The next steps are forecast validation by lifeguards and automation of feedback; these will allow the model to be continuously improved. As more data become available, other modeling strategies may be appropriate.

## Conclusions

Predicting the need for rescue from water in a hazardous environment is key to reducing the risk of drowning. Our predictive models can be used to efficiently deploy lifeguards and rescue helicopters. An interventional study (performed under real-world conditions) is planned. A utility function reflecting risk perception/acceptance is required. This would allow targeted preventative messages to be broadcast during high-risk periods. The strategy must employ behavioral change theory to reduce the risk to beachgoers. Evaluation requires reliable data from both lifeguard stations and emergency call center files.

The English in this document has been checked by at least two professional editors, both native speakers of English. For a certificate, please see: http://www.textcheck.com/certificate/N4bVZB

## References

1. Dicker D, Nguyen G, Abate D, et al. Global, regional, and national age-sex-specific mortality and life expectancy, 1950-2017: a systematic analysis for the Global Burden of Disease Study 2017. The Lancet. 2018;392(10159):1684–1735. doi:10.1016/S0140-6736(18)31891-9

2. Lasbeur L, Szego-Zguem E, Thélot B. Surveillance épidémiologique des noyades – Enquête NOYADES 2015. 1er juin – 30 septembre 2015. 2016. www.santepubliquefrance.fr.

3. Tellier É, Simonnet B, Gil-Jardine C, Castelle B, Bailhache M, Salmi L-R. Characteristics of drowning victims in a surf environment: a 6-year retrospective study in southwestern France. Inj Epidemiol. 2019;In Press. doi:10.1186/s40621-019-0195-x

4. Gensini VA, Ashley WS. An examination of rip current fatalities in the United States. Nat Hazards. 2010;54:159–175.

5. Bruneau N, Castelle B. Field observations of an evolving rip current on a meso-macrotidal well-developed inner bar and rip morphology. Cont Shelf Res. 2009;29:1650–1662.

6. Castelle B, Brander R, Tellier E, et al. Surf zone hazards and injuries on beaches in SW France. Nat Hazards. 2018;93(3):1317–1335. doi:10.1007/s11069-018-3354-4

7. Morgan D, Ozanne-Smith J, Triggs T. Descriptive epidemiology of drowning deaths in a surf beach swimmer and surfer population. Inj Prev. 2008;(14):62–65.

8. Li Z. Rip current hazards in South China headland beaches. Ocean Coast Manag. 2016;121:23–32. doi:10.1016/j.ocecoaman.2015.12.005

9. Arun Kumar SVV, Prasad KVSR. Rip current-related fatalities in India: a new predictive risk scale for forecasting rip currents. Nat Hazards. 2014;70(1):313–335. doi:10.1007/s11069-013-0812-x

10. Sherker S, Williamson A, Hatfield J, Brander R, Hayen A. Beachgoers’ beliefs and behaviours in relation to beach flags and rip currents. Accid Anal Prev. 2010;42(6):1785–1804. doi: http://dx.doi.org/10.1016/j.aap.2010.04.020

11. Bierens J. Handbook on Drowning. (Bierens JJLM, ed.). Berlin Heidelberg: Springer-Verlags; 2006.

12. Szpilman D, Bierens JJLM, Handley AJ, Orlowski JP. Drowning. N Engl J Med. 2012;366(22):2102–2110. doi:10.1056/NEJMra1013317

13. Lunetta P, Modell JH, Sajantila A. What is the incidence and significance of “dry-lungs” in bodies found in water? Am J Foren Med Pathol. 2004;25(4):291–301.

14. Hatfield J, Williamson A, Sherker S, Brander R, Hayen A. Development and evaluation of an intervention to reduce rip current related beach drowning. Accid Anal Prev. 2012;46(0):45–51. doi: http://dx.doi.org/10.1016/j.aap.2011.10.003

15. Szpilman D, Webber J, Quan L, et al. Creating a drowning chain of survival. Resuscitation. 2014;85(9):1149–1152. doi:10.1016/j.resuscitation.2014.05.034

16. Koon W, Rowhani-Rahbar A, Quan L. The ocean lifeguard drowning prevention paradigm: how and where do lifeguards intervene in the drowning process? Inj Prev. 2018;24(4):296–299. doi:10.1136/injuryprev-2017-042468

17. Szpilman D. Near-drowning and drowning classification: a proposal to stratify mortality based on the analysis of 1,831 cases. Chest. 1997;112(3):660–665.

18. Venema AM, Groothoff JW, Bierens JJLM. The role of bystanders during rescue and resuscitation of drowning victims. Resuscitation. 2010;81(4):434–439. doi:10.1016/j.resuscitation.2010.01.005

19. Canals M, Morell J. A nearshore breaker prediction system for Puerto Rico and the United States Virgin Islands in support of beach safety and drowning prevention. In: OCEANS’15 MTS/IEEE Washington. IEEE; 2015:1–10.

20. Cervantes O, Verduzco-Zapata G, Botero C, Olivos-Ortiz A, Chávez-Comparan JC, Galicia-Pérez M. Determination of risk to users by the spatial and temporal variation of rip currents on the beach of Santiago Bay, Manzanillo, Mexico: Beach hazards and safety strategy as tool for coastal zone management. Ocean Coast Manag. 2015;118:205–214. doi:10.1016/j.ocecoaman.2015.07.009

21. Austin MJ, Scott TM, Russell PE, Masselink G. Rip Current Prediction: Development, Validation, and Evaluation of an Operational Tool. J Coast Res. 2013;29(2):283–300.

22. Kerr KF, Meisner A, Thiessen-Philbrook H, Coca SG, Parikh CR. RiGoR: reporting guidelines to address common sources of bias in risk model development. Biomark Res. 2015;3(1). doi:10.1186/s40364-014-0027-7

23. von Elm E, Altman DG, Egger M, et al. The Strengthening the Reporting of Observational Studies in Epidemiology (STROBE) statement: guidelines for reporting observational studies. J Clin Epidemiol. 2008;61(4):344–349. doi:10.1016/j.jclinepi.2007.11.008

24. Beeck E van, Branche CM, Szpilman D, Modell JH, Bierens J. A new definition of drowning: towards documentation and prevention of global public health problem. Bull World Health Organ. 2005;83(11):801–880.

25. CEREMA Eau, mer et fleuves – ER/MMH. Candhis (Centre d’Archivage National de Données de Houle In-Situ). http://candhis.cetmef.developpement-durable.gouv.fr/.

26. Mazumdar M, Smith A, Bacik J. Methods for categorizing a prognostic variable in a multivariable setting. Stat Med. 2003;22(4):559–571. doi:10.1002/sim.1333

27. Heinze G, Wallisch C, Dunkler D. Variable selection – A review and recommendations for the practicing statistician. Biom J Biom Z. 2018;60(3):431–449. doi:10.1002/bimj.201700067

28. Hosmer DW, Hosmer T, Le Cessie S, Lemeshow S. A comparison of goodness-of-fit tests for the logistic regression model. Stat Med. 1997;16(9):965–980. doi:10.1002/(SICI)1097-0258(19970515)16:9<965::AID-SIM509>3.0.CO;2-O

29. Harrell, FE. Regression Modeling Strategies. Cham: Springer International Publishing; 2015. doi:10.1007/978-3-319-19425-7

30. DeLong ER, DeLong DM, Clarke-Pearson DL. Comparing the Areas under Two or More Correlated Receiver Operating Characteristic Curves: A Nonparametric Approach. Biometrics. 1988;44(3):837. doi:10.2307/2531595

31. Venkatraman E. A distribution-free procedure for comparing receiver operating characteristic curves for a paired experiment. Biometrika. 1996;83(4):835–848. doi:10.1093/biomet/83.4.835

32. Robin X, Turck N, Hainard A, et al. pROC: an open-source package for R and S+ to analyze and compare ROC curves. BMC Bioinformatics. 2011;12(1). doi:10.1186/1471-2105-12-77

33. R Core Team. R: A Language and Environment for Statistical Computing. Vienna, Austria: R Foundation for Statistical Computing; 2017. https://www.R-project.org/.

34. Fralick M, Denny CJ, Redelmeier DA. Drowning and the influence of hot weather. PLoS One. 2013;8(8):e71689. doi:10.1371/journal.pone.0071689

35. Clemens T, Tamim H, Rotondi M, Macpherson AK. A population based study of drowning in Canada. BMC Public Health. 2016;16:559. doi:10.1186/s12889-016-3221-8

